# Poplar: A Phylogenetics Pipeline

**DOI:** 10.1101/2024.11.11.623070

**Authors:** Elizabeth Koning, Raga Krishnakumar

## Abstract

**Motivation:** Generating phylogenetic trees from genomic data is essential in understanding biological systems. Each step of this complex process has received extensive attention in the literature, and has been significantly streamlined over the years. Given the volume of publicly available genetic data, obtaining genomes for a wide selection of known species is straightforward. However, analyzing that same data in order to generate a phylogenetic tree is a multi-step process with legitimate scientific and technical challenges, and often requires a significant input from a domain-area scientist.

**Results:** We present Poplar, a new, streamlined computational pipeline, to address the computational logistical issues that arise when constructing phylogenetic trees. It provides a framework that runs state-of-the-art software for essential steps in the phylogenetic pipeline, beginning from a genome with or without an annotation, and resulting in a species tree. Running Poplar requires no external databases. In the execution, it enables parallelism for execution for clusters and cloud computing. The trees generated by Poplar match closely with state-of-the-art published trees. The usage and performance of Poplar is far simpler and quicker than manually running a phylogenetic pipeline.

**Availability and Implementation:** Freely available on GitHub at https://github.com/sandialabs/poplar. Implemented using Python and supported on Linux.

**Supplementary Information:** Newick versions of the reference and generated trees.

## 1. Introduction

Creating phylogenetic trees from sequenced genomes is core to most areas of bioscience and biotechnology, as an understanding of the genomic relationships between organisms is critical for prevention, surveillance, intervention and optimization of biological processes (Kapli et al., 2020). In particular, understanding microorganismal diversity is critical both for defending against pathogens and for optimizing organisms to support the bioeconomy, with examples including bacteria, fungi and even members of the plant kingdom such as algae (Spatafora et al., 2017; Nakano et al., 2023; Navarro, 2021; Azouri et al., 2021). In addition, phylogenetics play a critical role in viral, bacterial and fungal pathogen and pandemic monitoring, as exemplified recently by SARS CoV2 and fungal plant pathogens (De Maio et al., 2023; Turakhia et al., 2022; Ki and Terhorst, 2022; Dort et al., 2023).

To date, there are numerous strategies and methods associated with building these trees. During this process, it is essential to combine biological meaning with computational efficiency and flexibility. While early approaches relied on observable physical and metabolic characteristics, the expansion of genetics and genomics over the decades has added a significant amount of resolution to these methods (Hug et al., 2016b).

Building a species tree requires genomic data from each included species to be sequenced, analyzed, compared, and then constructed into a tree. The specific steps may differ, but a typical pipeline includes sequencing, identifying homologs, aligning sequences, inferring gene trees, and finally constructing a species tree (Young and Gillung, 2020). Each step requires one or more software tools requiring various inputs and data formats. In addition, software that offers sequence-to-tree analysis has its own limitations.

For example, MEGA is a comprehensive tool for tree-building (Tamura et al., 2021). However, MEGA requires either a genome annotation or whole genome multiple genome alignment, and while this can be extremely helpful for some applications, it can have limitations in other situations, for example, if users want to focus on specific gene subsets without requiring significant manual intervention, which might be challenging for non-subject matter experts. Another challenge a user may face if attempting to use whole genomes is the requirement of a single sequence for each species, which excludes assemblies of scaffolds. If there is an annotation, the user must use another tool to group gene sequences before alignment. Furthermore, MEGA is not open source, so users cannot adapt it or run on incompatible operating systems. It also does not support parallelism.

EasyCGTree is another tool for inferring phylogenetic trees Zhang et al. (2023). It includes options for multiple inference methods and does include a step to group sequences to form gene trees. However, it does not include parallelism, which is a significant roadblock when building increasingly larger trees.

A third software tool for tree inference is Read2Tree (Dylus et al., 2023). While the authors of Read2Tree report highly accurate output trees, Read2Tree requires an internet connection during its run as well as a download from the author’s Orthologous Matrix (OMA) database before running. For some applications, the required remote connection would be a security concern or a practical restriction. As for the data required from the external database, the database does not include sequences for all species. While we have not done a comprehensive examination of the included species, in our comparative testing, we found OMA includes very few fungal sequences. Read2Tree also uses reads, rather than assemblies. In some cases, this may be very useful, but if genomes are selected from the NCBI database, they will typically have assemblies available, which allows for downloading and managing a much smaller amount of data. In addition, reads can be assembled into genomes or transcriptomes, but the reverse is not true.

To address the issues of ease of use, parallelism and flexibility outlined above, we present Poplar, a novel software tool for the pipeline from assembled genome to phylogenetic tree. As a tree species, “poplar” connects the concept of biological trees with the phylogenetic trees the software infers. Specifically, poplar trees have a reputation for fast growth, and Poplar, the software tool, is designed to quickly grow a phylogenetic tree (Britannica, 2023). Tasks as seemingly simple as converting file types or as time consuming as genome annotation can bog down the development of a phylogenetic inference pipeline, and Poplar addresses the issue by smoothing the connections between the numerous steps in tree inference and using annotated or unannotated genomes.

## 2. Background

To obtain a species tree from a collection of assembled genomes typically requires a variety of software packages. While each step in the pipeline has well established and ever improving tools, those tools have unique file formats and input requirements that are frequently incompatible with other tools. Many steps have entirely independent computation, but executing in parallel requires manually initiating parallel jobs. Poplar addresses these concerns and manages converting file types, renaming sequences, and running tasks in parallel.

The key steps are:

1. Assemble genome from reads (if using next-generation sequencing data as opposed to a pre-assembled genome)
2. Identify gene sequences
3. Group sequences into genes across species
4. Align gene sequences
5. Create gene trees
6. Merge gene trees into species tree

There may be variation in these steps, many of which are described in (Kapli et al., 2020). Some approaches will require whole genome alignment (WGA) rather than perform alignments for gene sequences, for example. WGA is done through specialized tools such as Cactus (Armstrong et al., 2020) or more general alignment tools such as RAxML-NG and Muscle (Kozlov et al., 2019; Edgar, 2022). WGA is a significantly costly computation and therefore larger alignments are often not feasible on computational resources that are readily available to most researchers. Kmer-based distance methods can also substitute for alignment, and they bypass using full gene sequences at all, but rather use dictionaries of kmers. These methods are extremely helpful for fast examination of relationships between genomes, particularly large numbers of large genomes Nasko et al. (2018); Manekar and Sathe (2018); Moeckel et al. (2024). However, the caveat of using kmer-based methods is that the gain in speed sacrifices some accuracy, and while this loss in accuracy can be mitigated by having more individual genomes to analyze (i.e. more pairwise distance comparisons can make for a more accurate overall tree), we focused Poplar to use methods (discussed below) that are more resilient and accurate even with low genome/sequence numbers.

Within gene-driven trees are a specific subset that are frequently used - alignment based on ribosomal genes. While there are numerous examples of this in the literature, a significant one is the recent novel view of the tree of life, where multiple ribosomal genes were used (Hug et al., 2016a). While focusing on ribosomal genes can standardize comparisons, it can be limiting in cases needing discernment of similar organisms, or strains within a species, where the ribosomal genes are likely to be very similar and may not provide the desired resolution.

Most often, phylogenetic trees are created through ad-hoc pipelines, such as in the 1000 Plants Initiative and the Avian Phylogenomics Project Wickett et al. (2014); Jarvis et al. (2015). This has the benefit of allowing the scientists to use tools they expect to be most suitable for their data in each step, but it also requires implementing many elements of the application, such as the tools to connect the main steps. Ad hoc pipelines also face challenges in presentation. The methods to construct published trees are often not specific enough for other researchers to adapt or reproduce. Since code for the pipelines are often not included with publications, each research group must implement their own pipeline, which may or may not share characteristics with other pipelines, leading to wasted energy and a lack of clarity in the origins of various phylogenetic trees. In designing Poplar, our goal is to implement a tried-and-tested combination of the algorithms required to complete the key steps outlined below. We acknowledge that there are alternatives, and ultimately the long-term goal of Poplar is to be modular and allow for flexibility in algorithm usage, but we describe here a pipeline that uses established tools in an efficient and accessible way. Specifically we focus on optimizing a few processes outlined below.

Inferring gene trees requires collections of gene sequences. Identifying genes as genes that occur across multiple species requires a genome annotation. De novo gene prediction identifies probable genes based on patterns within the genomic sequence, while alignment based methods rely on a reference database in order to compare sequences in a genome with known genes (Vallender, 2009). Either approach is a significant task in terms of both labor and computation. On the other hand, identifying homologs that do not have an identified function is a simpler task but provides far less information. They may resemble one another due to shared ancestry or horizontal gene transfer, or the sequences may have no relation. Occasionally, this task is done with grouping algorithms such as DBSCAN (Edla and Jana, 2012). If an annotation is available for some genomes of interest but not others, that typically means that the unannotated genomes will be used for all included species.

As described by (Kapli et al., 2020), gene tree inference may be done through techniques including distance methods, such Neighbor Joining (NJ), or character-based methods, such as Maximum Likelihood (ML). Deciding which method is most practical for a particular situation is dependent on factors such as how closely related the sequences are.

Given a set of gene trees, various supertree methods also exist to infer the species trees. A popular tool for species tree inference is ASTRAL-Pro3 (Zhang and Mirarab, 2022), which optimizes for the number of quartets occurring in the gene trees in the resulting species tree. Poplar incorporates ASTRAL-Pro3 as a method for fusing the gene trees generated in the previous step.

## 3. Algorithm

Poplar provides a structure to connect established tools for each step of phylogenetic tree inference. Beginning from gene or genome data, it works through identifying relevant sequence, building possible gene groupings, inferring gene trees, and constructing a species tree. Figure 1 illustrates the pipeline.

**Fig. 1:**
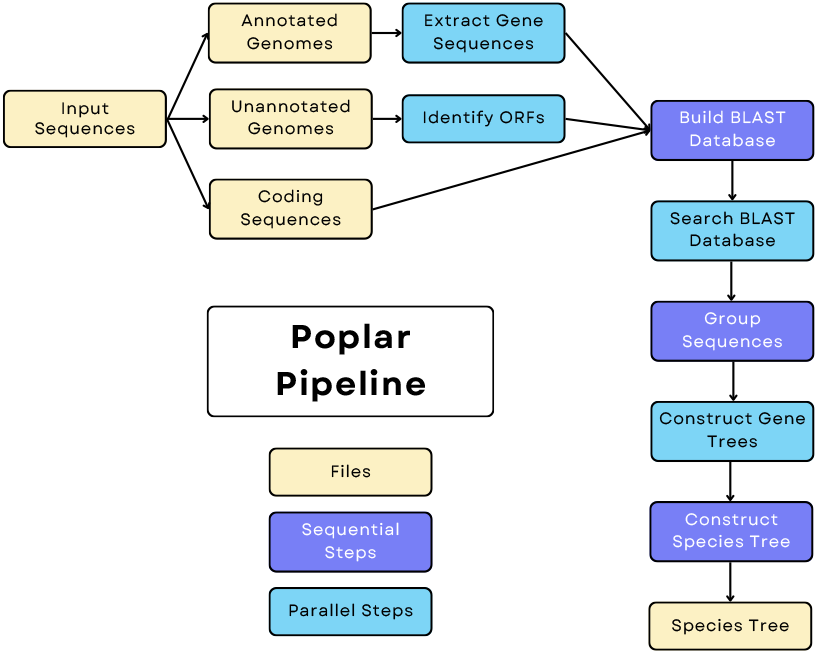
Illustration of the Poplar Pipeline

Poplar does not have a next-generation sequencing analysis or genome assembly module, and assumes you are providing either genomes, scaffolds or genes. We will describe the other key steps identified in the background section which are implemented by Poplar.

### 3.1. Identification of gene sequences

The first step in Poplar is identifying gene sequences for each input species. For some species, these will be input as sequences, and for others, the sequences must be extracted from the genome scaffold sequences. If coding sequences are provided for a species, those sequences will be prioritized. Otherwise, if an annotated genome with identified genes is provided, then the sequences are extracted and used as potential genes. If not, then Open Reading Frames (ORFs) are extracted from the genome using orfipy (Singh and Wurtele, 2021). In a single run of Poplar, each type of data may be used for different genomes. Currently, the implementation requires nucleotide sequences.

### 3.2. Sequence Grouping into Genes Across Species, and Alignment of Gene Sequnces

Following the identification of genes and ORFs, Poplar groups the sequences in clusters.

First, Poplar uses Nucleotide BLAST to identify similar sequences across genomes. All the sequences are collected in a Nucleotide BLAST database Altschul et al. (1990). Poplar then queries the database for each sequence. The default similarity threshold is an expectation value (evalue) of 1^−20^, but this threshold may be increased or decreased by the user if the input organisms are less or more closely related, respectively.

The results of the searches are used to create a distance matrix, which is then fed into an implementation of Density-Based Spatial Clustering of Applications with Noise (DBSCAN) for grouping (Ester et al., 1996; Schubert et al., 2017). DBSCAN was selected for this step because it does not force a fixed number of groups, a fixed size of groups, or all sequences to be placed into a group. Instead, it identifies dense clusters as sequences that it declares groups. After using DBSCAN to create these groups, Poplar omits groups that are too small or too large, based on user input. By default, the minimum group size is 3, and the maximum group size is 100.

Using a grouping algorithm like DBSCAN to identify homologs across identified gene sequences is not the most popular way that gene clusters are identified. Often, orthologs will be identified and stored in a database such as OMA for future reference (Vallender, 2009). Using the BLAST search followed by DBSCAN only takes into account the similarity between two sequences, not any contextual information about their respective genomes. Yet, DBSCAN has been used for identifying gene clusters, including as an aspect of the tool GeneGrouper (McFarland et al., 2021). The results of Poplar also demonstrate the effectiveness of using a distance-based clustering for sequences.

Rather than rely on an external database or an intensive annotation process, Poplar considers a large set of sequences, and identifies those that are most similar. The BLAST database is stored locally, allowing Poplar to execute offline. The pairing of BLAST and DBSCAN allows Poplar to identify similarities between sequences, including sequences from genomes that have not been annotated.

The output of the sequence grouping step is the sets of sequences identified as similar.

### 3.3. Creation of Gene Trees

Once Poplar has a set of sequence groups, it selects a subset of the groups to build gene trees. By default, it will randomly select 50 groups, and a user may specify some other maximum number of trees to build.

Poplar infers gene trees from the sequence groups via MAFFT (Katoh and Standley, 2013) for sequence alignment and RAxML-NG (Kozlov et al., 2019) for gene tree construction. By default, Poplar also uses the default settings for both tools. The user may select the maximum number of gene trees to create, as the number of groups available to create trees is likely to be high, due to the nature of the sequence grouping process. MAFFT and RAxML-NG are both established tools in the field. RAxML-NG outputs the best tree for each set of sequences, which is then the input for creating the species tree.

### 3.4. Merging of Gene Trees into Species Trees

In order to infer a species tree from a set of gene trees, Poplar uses ASTRAL-Pro3 (Zhang and Mirarab, 2022). Like MAFFT and RAxML-NG, ASTRAL is an established and popular tool. The output of ASTRAL is a single tree including each of the input species.

## 4. Implementation

Poplar works as a framework or backbone to connect various well established tools. It runs as a Python script that manages inputs and refers to other tools, allowing the user to select options such as the number of gene trees, the maximum number of sequences to include in a gene group, and the evalue threshold for BLAST. In order to manage parallelism, it uses Parsl (Babuji et al., 2019), a Python workflow library. Parsl allows Poplar to fluctuate the hardware allocated to the tasks as needed.

Our tests were performed on a system using Slurm, but Parsl also supports running a workflow locally, or via other schedulers or cloud services. Changing the type of system used is done in a configuration file, and a Parsl “provider” is selected. The configuration file must be customized for the system being used in order for Parsl use the correct Slurm queue or other method of accessing the hardware.

Some steps required new code within Poplar. Most of the new code involves changing file formats and data labels, and are clearly outlined in our user manual. The one more substantial element is in identifying homolog groups using DBSCAN, which are then used for the “gene” trees. The algorithm cannot confirm that the groups represent a single gene across the species. Rather, it merely groups similar sequences together. However, as we will demonstrate in the results, and based on prior literature, this is an effective method for finding groups of sequences that can be annotated as individual genes for tree-building (McFarland et al., 2021).

Not using new, untested algorithms is a strength of the current iteration of Poplar –because it does not utilize new ways of generating gene trees or species trees, the end result when using Poplar is the same as when using RAxML or ASTRAL-Pro, thereby giving expected and trusted results in a more streamlined and efficient way. When new software is introduced or different tools and models are needed for particular uses, Poplar can also be updated to use the most accurate algorithms. The accuracy is that of its component tools, and has the same theoretical and applied accuracy as its components. A user experienced with the component tools may want to modify the calls to these tools to use alternative options from the default configuration of each piece of software.

The input to Poplar is FASTA files. As discussed earlier, these may be coding sequences, annotated genomes, or unannotated genomes. The intermediate data from each of these steps may be preserved for user reference or be deleted to minimize stored files. Poplar also supports check-pointing through Parsl. The clustering algorithm outputs lists of sequence identifier for all groups it creates. MAFFT outputs an alignment file in FASTA format, and RAxML outputs gene trees in Newick format. The output of Poplar is a Newick tree, including branch lengths.

## 5. Results

### 5.1. Testing Data

The testing data came from NCBI, JGI’s MycoCosm, and JGI’s Phycocosm. Specifically, fungal genomes were chosen due to the fact the fungal kingdom is both highly diverse and rapidly growing, and the use of phlogenetic trees to characterize new and emerging fungal species is key and widespread (Frac et al., 2018; Shumskaya et al., 2023; Naranjo-Ortiz and Gabaldón, 2019). The test datasets are:

1. The 13 Kickxellomycotina genomes available through MycoCosm (Reynolds et al., 2023; Chang et al., 2015; Amses et al., 2022; Mondo et al., 2017; Wang et al., 2016; Ahrendt et al., 2018)
2. The 88 Mortierellomycotina genomes available through MycoCosm (Vandepol et al., 2020; Mondo et al., 2017; Mathieu et al., 2024; Chang et al., 2022; Mesny et al., 2021; Uehling et al., 2017)
3. The 26 Rhodophyla coding region sets available through PhycoCosm (Schönknecht et al., 2013; Rossoni et al., 2019; Cho et al., 2023; Nozaki et al., 2007; Lee et al., 2018; Nakamura-Gouvea et al., 2022; Collén et al., 2013; Brawley et al., 2017; Nakamura et al., 2013; Bhattacharya et al., 2013)
4. NCBI’s 20 commonly used organisms for molecular research projects (Assembly, 1988a,b,c,d,e,f,g,h,i,j,k,l,m,n,o,p,q,r,s,t)
5. The 5 genomes of Puccinia graminis available through MycoCosm Duplessis et al. (2011); Li et al. (2019)

For the datasets from JGI, the reference trees came from the JGI webpage. For the NCBI dataset, the reference tree is from the species classifications. The reference trees do not include branch lengths.

A key criteria in selecting the datasets was the availability of a fully resolved reference tree from clear and well-grounded methodology. JGI publicly describes their Annotation Pipeline, and all included genomes are presented in the species tree, including where there are multiple genomes from a single species. NCBI presents only the classification of species, which means the tree has many polytomies. Polytomies are not an issue with the 20 common species as they are distantly related but are an issue with any closely related set of species or any species with multiple variants.

### 5.2. Metrics

We report the accuracy of trees by the Robinson-Foulds distance. The Robinson-Foulds metric works by identifying all bifurcations of the trees, and comparing the bifurcations in one tree to those of the other. If a split is present in only one of the two trees, the distance score increases by one. The reported distances are normalized to be out of 100%, where 0% is an identical tree and 100% shares no bifurcations.

Where the reference tree is not a binary tree, we do not report the additional splits in the generated trees as inaccuracies. MycoCosm presents binary trees, while NCBI provides the classifications of the species, which are not binary trees but also do not assert that there are not additional relationships that are not captured by the classification schema. Poplar reports binary trees.

### 5.3. Kickxellomycotina

The Kickxellomycotina classification within fungi has 13 genomes in JGI’s MycoCosm, with the tree shown in Figure 2. Some species within Kickxellomycotina have multiple genomes represented. For this test, we used the genome assemblies rather than the annotated genomes, even when an annotation was available. When run five times with 50 random gene trees generated, the average Robinson-Foulds error rate is 0.20. The results of one run with a 11% Robinson-Foulds error rate is shown in Figure 3. As shown in Table 1, the average error with 100 gene trees is 11% and with 500 gene trees is 7%.

**Table 1.** Poplar Performance results on Kickxellomycotina unannotated genomes.

| Input Type | Number of Gene Trees | Number of Runs | Average RF Error Rate | Minimum RF Error Rate | Maximum RF Error Rate |
| --- | --- | --- | --- | --- | --- |
| Unannotated genome | 50 | 5 | 0.20 | 0.11 | 0.33 |
| Unannotated genome | 100 | 5 | 0.11 | 0 | 0.22 |
| Unannotated genome | 500 | 5 | 0.07 | 0 | 0.22 |

**Fig. 2:**
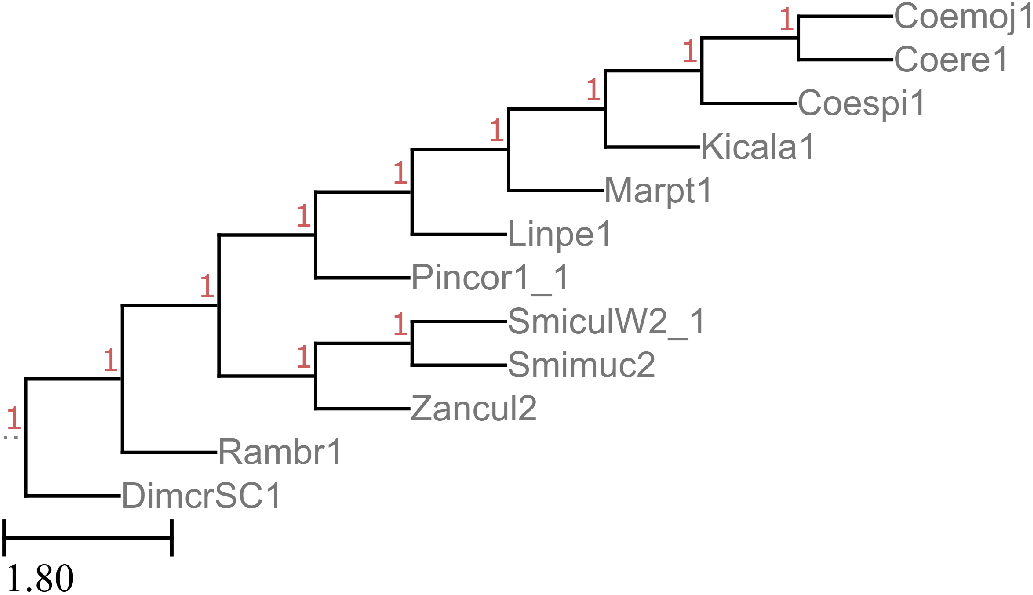
Tree of Kickxellomycotina genomes, according to JGI’s MycoCosm

**Fig. 3:**
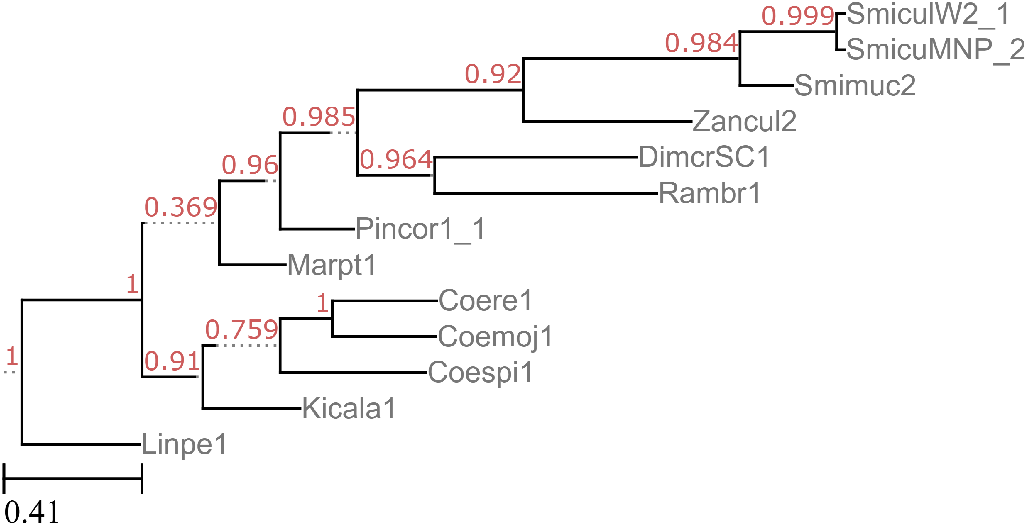
Tree of Kickxellomycotina genomes, according to one run of Poplar with 50 gene trees

### 5.4. Mortierellomycotina

Mortierellomycotina is another fungal classification, with 88 genomes in JGI’s MycoCosm. As with Kickxellomycotina, some species have multiple genomes in the database, and the tree was generated from genome assemblies, omitting annotations. In a run with 200 gene trees, Poplar’s tree differed from the MycoCosm tree by 21 out of 86 splits, having a Robinson-Foulds distance of 24%. The Newick versions of the trees are included in the supplementary materials.

### 5.5. Rhodophyla

Rhodophyla is a algal classification, with 26 genomes available through JGI’s PhycoCosm, 20 of which are included in their tree. In a single run with 200 gene trees, Poplar had a distance of 15% from PhycoCosm’s tree. The two trees are shown in Figure 4 and Figure 5.

**Fig. 4:**
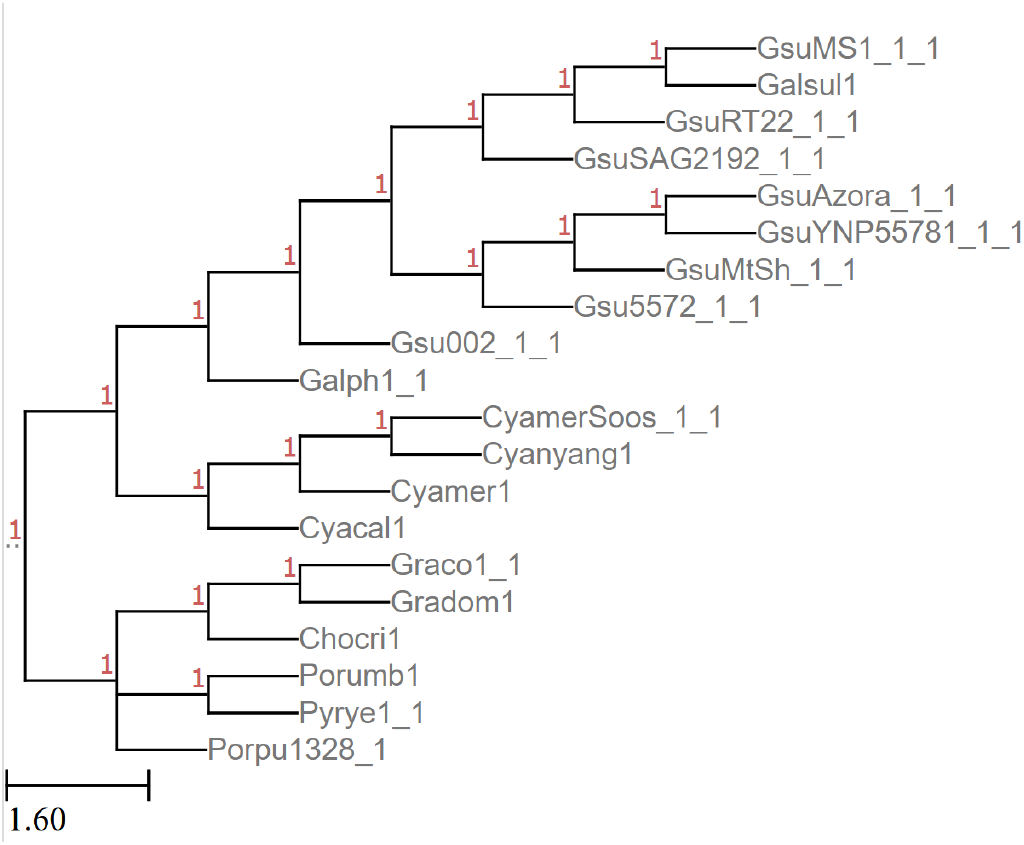
JGI’s Rhodophyla Tree

**Fig. 5:**
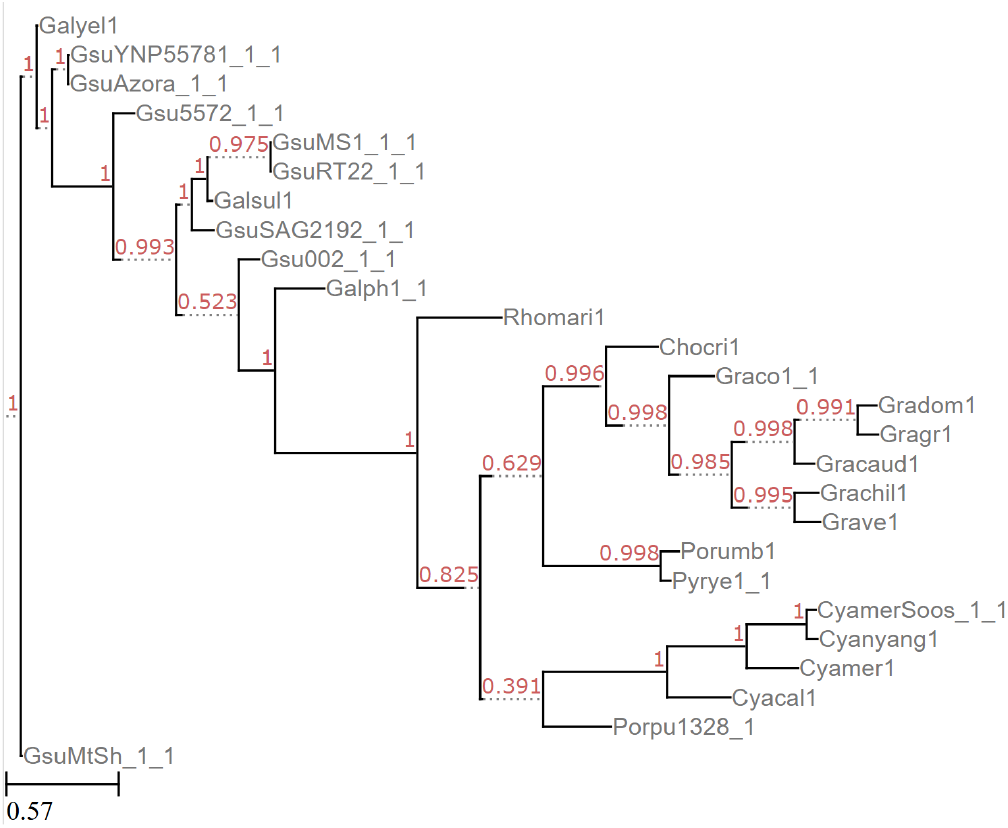
Rhodophyla tree generated by Poplar with 200 gene trees

### 5.6. NCBI Commonly Studied Organisms

These 20 organisms from NCBI’s database were included in order to demonstrate the usage of Poplar on genomes outside of fungi and algae, as well as on a tree with a vast spread of species.

Two runs of Poplar, both with 200 gene trees, resulted in slightly different trees. Because of the random selection of the groups, Poplar is not deterministic. Given enough gene trees, the accuracy impact of the randomness will typically be small, while the runtime impact may be significant.

The output of Poplar did not include *Mycoplasmoides pneumoniae* or *Escherichia coli*. This exclusion is likely due to their distance from the other species. The other difference was in the placement of *Plasmodium falciparum*. In the first run, it was placed among the plants, leading to a Robinson-Foulds distance of 20%. In the second run, it was placed accurately next to *Dictyostelium discoideum*, and therefore the Robinson-Foulds distance was 0%. The misplacement of *P. falciparum* was most likely due to the large spread of the included species. If a species does not share as much genetic material with the other species in the tree, it will be included in fewer groups and gene trees, and a misplacement in one tree that might otherwise be noise can be perpetuated into the species tree.

### 5.7. Puccinia graminis

The five genomes of the species Puccinia graminis are included to contrast the diversity of the NCBI Commonly Studied Organisms, and demonstrate the effectiveness of Poplar on genomes that are closely related. Error rates are shown in Table 2. Runs with 50 gene trees varied between each other, though the distance from the MycoCosm tree was consistent for all trees. Using 100 or 500 gene trees, all five trees were the same as each other, though they varied from MycoCosm’s tree. As shown in Figure 8, the two trees vary in therelationship between Pucgr2 and Pgt 201 B1. The node in MycoCosm’s tree connecting (Pgt Ug99 A1, Pgt 201 A1) to Pgt 201 B1 has much lower support than the other nodes in the tree. While all other nodes have the maximum support of 1.0, this split has support of only 0.2850. It is plausible that the difference between the MycoCosm and Poplar trees is supported by the available data.

**Table 2.** Poplar Performance results on Puccinia graminis coding sequences.

| Input Type | Number of Gene Trees | Number of Runs | Average RF Error Rate | Minimum RF Error Rate | Maximum RF Error Rate |
| --- | --- | --- | --- | --- | --- |
| CDS | 50 | 5 | 0.50 | 0.50 | 0.50 |
| CDS | 100 | 5 | 0.50 | 0.50 | 0.50 |
| CDS | 500 | 5 | 0.50 | 0.50 | 0.50 |

**Fig. 6:**
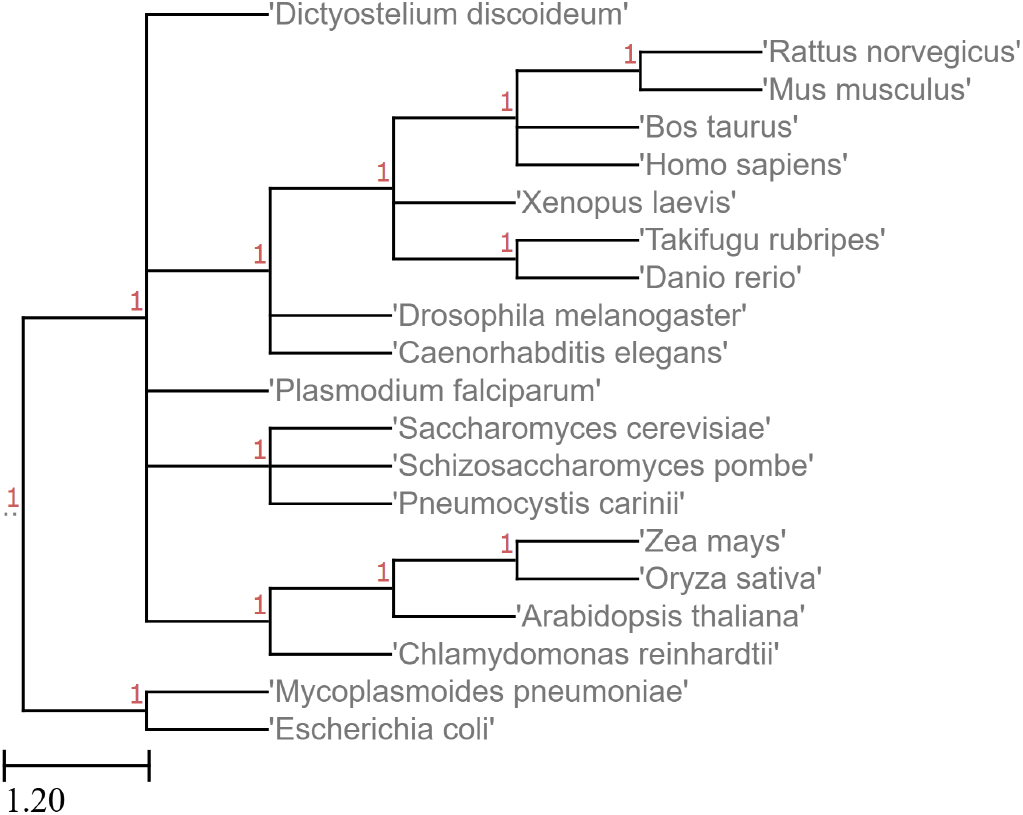
True tree of species from NCBI

**Fig. 7:**
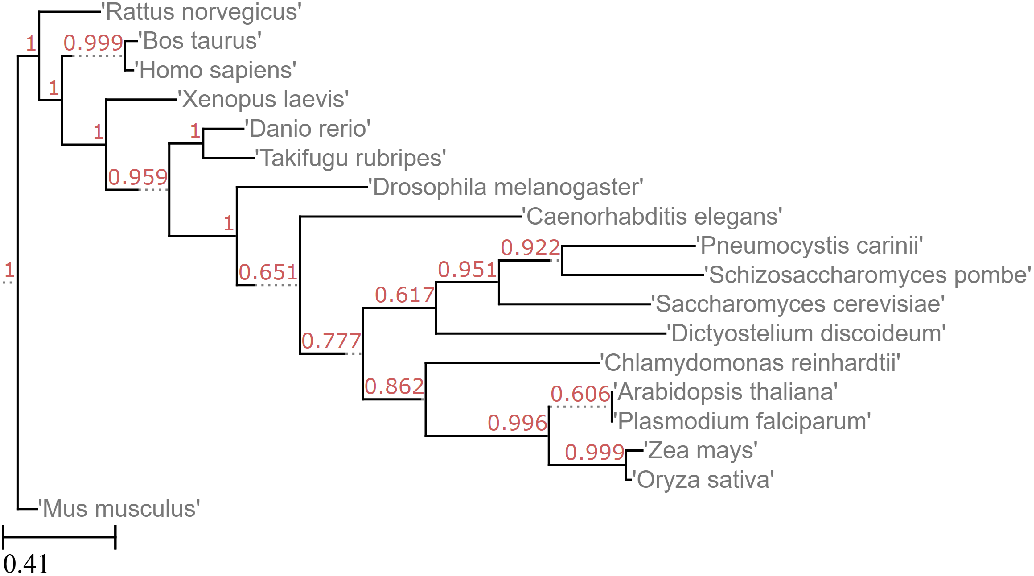
Tree generated by Poplar from the NCBI genomes

**Fig. 8:**
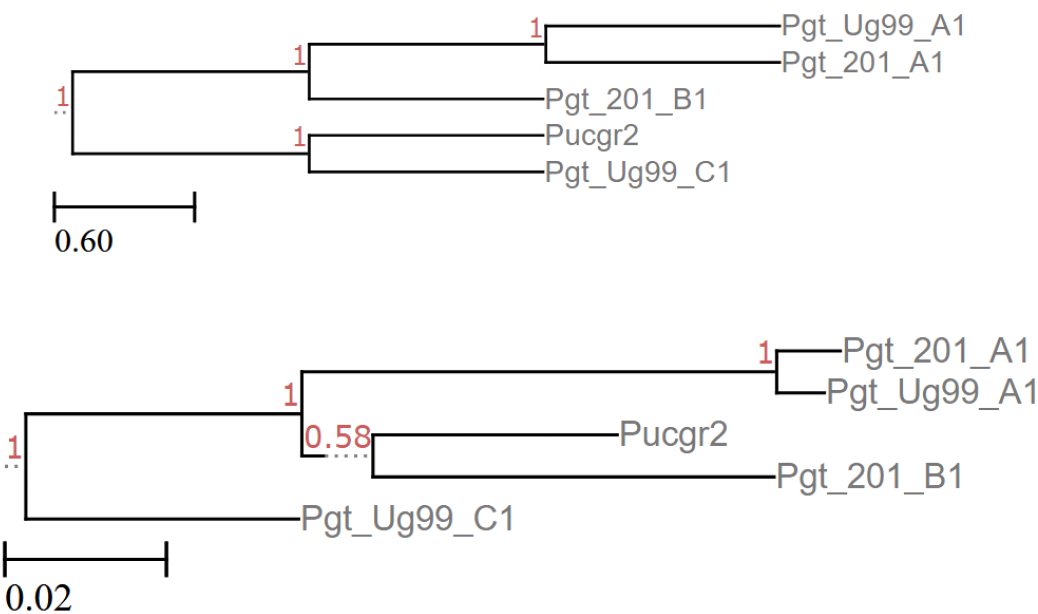
Puccinia graminis trees from MycoCosm (top) and Poplar with 100 or 500 gene trees (bottom)

### 5.8. Performance

We assessed performance on the 14 Kickxellomycotina genomes, using the full genomes. The tests were run on CPU nodes on a High Performance Data Analytics Research Cluster with Dual Socket Intel E5-2683v3 2.00GHz CPUs.

In these runs, each worker uses 2 cores, so each node is able to run 28 tasks at a time. Two nodes were used per block, and Parsl was permitted a maximum of 4 blocks.

Table 3 shows the performance of Poplar when running 100 trees. The majority of the time is spent in building the gene trees. While the total time is nearly 12 hours, the runtime from start to end was actually only 23 minutes. The division of each node into 28 workers also served to reduce the node usage.

**Table 3.** Time per task in building the Kickxellomycotina tree with 100 gene trees.

| Task | Number of Invocations | Average Time | Total Time (hh:mm:ss) |
| --- | --- | --- | --- |
| Find ORFs | 14 | 00:00:11 | 0:02:39 |
| Build BLAST Database | 1 |  | 0:00:09 |
| Search BLAST | 14 | 0:02:33 | 0:35:52 |
| Make groups | 1 |  | 0:05:30 |
| Align Sequences and Construct Gene Trees | 100 | 0:06:36 | 11:00:44 |
| Other | 6 | 0:01:51 | 0:11:08 |
| ASTRAL-Pro | 1 |  | 0:00:00.37 |
| Total |  |  | 11:56:04 |

In the same way, Table 4 shows the performance of Poplar when running 500 trees for Kickxellomycotina. Again, generating the gene trees was where the majority of time was spent.

**Table 4.**
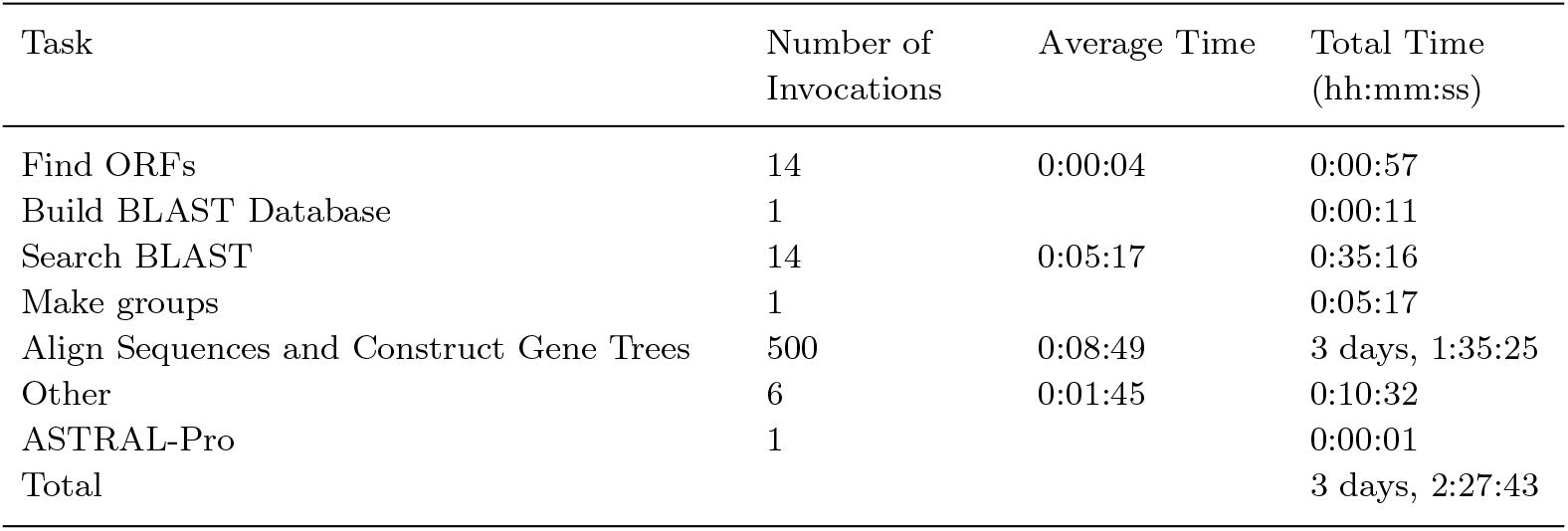
Time per task in building the Kickxellomycotina tree with 500 gene trees.

Figure 9 shows the CPU usage throughout the run with 500 trees. During significant portion of the run, the parallelism is able to use all available CPUs.

**Fig. 9:**
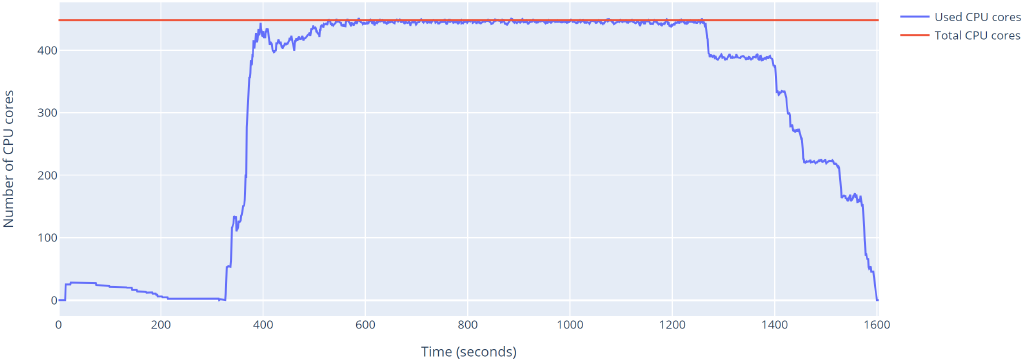
Graph of CPU usage during execution of Poplar on 14 Kickxellomycotina genomes using 500 gene trees

## 6. Discussion

Poplar offers a readily usable alternative to building custom phylogenetic tree pipelines, which often require subject-matter expertise, and the ability to string together multiple pieces of software that do not necessarily have compatible inputs and outputs. In addition, Poplar can utilize genomes with or without annotation, and advanced users can vary the number of gene trees that get constructed, thereby increasing the resolution of the tree. This higher-resolution tree will require more resources, giving users the option to choose between high resolution and speed/resource utilization.

The implementation of Poplar allows flexibility across hardware. While testing was only performed on clusters using Slurm, Parsl supports cloud computing and other cluster management systems. Shifting between systems across workflows or within a single workflow can be done in a configuration file, with Parsl offering examples and support for a wide variety of architectures. While Poplar is limited by the operating systems and hardware where the other software is depends on is able to run, some steps are portable to non-Unix machines and could be ported more widely with minimal user intervention.

Poplar’s results match closely with reference trees from accepted sources. Each step in Poplar is not new. Instead, the algorithms and software it incorporates are widely used and based on substantial theoretical and practical study. The novel aspect of Poplar is in the ways it allows a user to more easily run the entire pipeline with minimal interference while still obtaining results from state-of-the-art algorithms. Gathering the phylogenetic inference pipeline into one tool allows for coherency of use and communication.

### 6.1. Limitations and Future Work

Poplar is designed in a modular manner, so a user may replace a step with an alternative tool. Using an alternative tool will also require providing scripts to convert input or output files if they differ from the default. Future work will include options for different methods for each step, allowing the change to be done by simply changing an argument. Adding additional options for each step will allow users who want to use particular tools or models to simply select the suitable tools via the command line or a configuration file.

Most runs of Poplar are dominated by searching the custom BLAST database for similar sequences. The time spent in this step is especially pronounced when the included genomes do not have provided coding regions or annotations. When sequence selection is not narrowed to only the coding regions or annotated genes, all ORFs in the genome are included, ballooning both the search space and the number of queries. Future work will look into both narrowing the number of queries used as well as more efficiently searching the database, given the goal of obtaining clusters.

Another area of future development of Poplar improving results with distantly related species. When the sequence groups are selected, the currently algorithm selects a random set of groups. When some species are distantly related from the main group, they may not have sequences occur in any of the groups. In that case, they will be omitted from the species tree. In the future, the selection of sequence groups will include a optimization for including all species.

## 7. Availability and requirements

Poplar is available at https://github.com/sandialabs/poplar under the GPLv3 license.

## Acknowledgments

We would like to thank Ellis Torrance and Sharon Nademanee for their review of the manuscript. This work was supported by the Laboratory Directed Research and Development program at Sandia National Laboratories, a multi-mision laboratory managed and operated by National Technology and Engineering Solutions of Sandia, LLC, a wholly owned subsidiary of Honeywell International, Inc., for the U.S. Department of Energy’s National Nuclear Security Administration under contract DE-NA0003525.

